# Antiviral Potential of Lauric Acid against Dengue Virus 2: Evidence from a Luciferase-Based Replicon Assay

**DOI:** 10.64898/2026.04.23.720334

**Authors:** Anjali Kumari, Rajendra Pilankatta, Bharti Kumari, Mihir Kumar Prasad, Nikhil Kumar, Anubha Kumari

## Abstract

Dengue virus (DENV) infection generates a significant health burden throughout the world, and there are no clinically approved antiviral drugs, as of now. The virus also depends on lipid metabolism in the host to conduct effective replication and this makes lipid-directed compounds promising as therapeutic options. We assessed the antiviral effect of lauric acid, a 12-carbon medium-chain fatty acid, against DENV serotype 2 (DV2) in the presence of a stable cell line, DV2-replicon, expressing all the non-structural proteins (NS1-NS5) and a luciferase reporter. Active viral replication in replicon cells was established by morphological examination and immunofluorescence of cells. The MTT assay was used to determine the cytotoxicity of lauric acid revealing the LD50 of 2.52 uM, so higher concentrations were toxic as the effect of the drug is dose-related. The antiviral effect was tested through replicon inhibition (luciferase) assay which showed an incredible inhibition of viral RNA replication with a IC50 of 1.70 uM and this is equivalent to antiviral mycophenolphycic acid. The cytopathic effects, as well as a decrease in the activity of luciferase, proved the presence of viral translation and replication inhibition within the process of the treatment of the lauric acid. These results propose that lauric acid has cytotoxic and antiviral dual effect and can be a possible inhibitor of DENV replication. The toxicity needs to be reduced and future research is necessary to explain its molecular pathway and also to come up with the best delivery methods.

**IMPORTANCE:** Dengue virus (DENV) remains a significant health challenge to the world since there are no effective antiviral agents. This work will recognize lauric acid as a possible dengue virus replication inhibitor in a model of a DV2 replicon, exhibiting antiviral action that is similar to that of mycophenolic acid. These results support lipid-directed compounds as potential dengue antiviral targets, but more research is needed to minimize toxicity and better understand the molecular mechanism of action.

## INTRODUCTION

Dengue is a viral infection spread through mosquitoes, mainly Aedes albopictus and Aedes aegypti (1). Dengue virus belongs to the genus Flavivirus in the family Flaviviridae, which also includes other serious viruses responsible for diseases such as Japanese encephalitis, West Nile virus, yellow fever, and tick-borne encephalitis (2). The disease caused by dengue can vary from a mild fever to a more serious condition called severe dengue, earlier referred to as dengue hemorrhagic fever (DHF). This severe form is marked by issues in blood clotting and leakage of fluids from blood vessels, resulting in low blood volume, thickening of the blood, and low blood pressure—symptoms that can turn fatal (4). Severe dengue was mainly reported during outbreaks in the Philippines and Thailand around 1950 (3). Currently, the disease is widespread in Asia and Latin America, where it is a major reason for hospital admissions and deaths among both children and adults.

### 1.1. Global Statistics of Infection of Dengue Virus

Dengue poses a threat to an estimated 2.5 billion individuals and has become an increasingly significant public health concern. Annually, dengue infects approximately fifty to one hundred million people, with close to 500,000 cases necessitating hospital admission because of severe and potentially life-threatening conditions. Both the overall incidence of dengue and the frequency of large-scale outbreaks have risen sharply in recent years. Earlier assessments showed that globally there are between 50 and 100 million cases of dengue fever, along with about 500,000 instances of dengue hemorrhagic fever, leading to roughly 22,000 fatalities, predominantly affecting children (5). An estimated 2.5 to 3 billion people—about 40% to 50% of the global population—across 112 tropical and subtropical countries is considered at risk for dengue infection (6). Antarctica remains the only continent where dengue transmission has not been observed (7).

**Table 1.** Association of Dengue Serotypes with Disease Severity.

| Dengue Serotype | Association to Dengue disease |
| --- | --- |
| DV-1 | Primary infection with this serotype often causes more severe illness compared to infections caused by DV-2 or DV-4 |
| DV-2 | Secondary infection with this serotype is linked to a higher risk of severe disease, with the likelihood of developing dengue hemorrhagic fever being nearly twice that seen with DV-4 |
| DV-3 | This serotype tends to cause more severe illness during a primary infection compared to DV-2 or DV-4. Additionally, during a secondary infection, it is nearly twice as likely to lead to dengue hemorrhagic fever compared to DV-4 |
| DV-4 | This serotype is the least likely to cause severe dengue complications |

A positive-sense single-stranded RNA ((+) ssRNA) virus has RNA that functions directly as messenger RNA (mRNA) upon entering the host cell, allowing its genetic code to be immediately translated into proteins. RNA viruses are categorized according to the polarity of their RNA strands, being either positive-sense or negative-sense. For positive-sense RNA viruses, the host’s cellular machinery can translate their genomes right away. Examples of viruses in this group include dengue virus, hepatitis C virus, West Nile virus, coronaviruses responsible for SARS and MERS, and common cold viruses such as rhinoviruses (8).

### 1.2. Genome Organisation of Dengue Virus

The dengue virus is an enveloped virus containing a positive-sense, single-stranded RNA genome. Its genetic material can be directly translated into proteins once inside the host cell. The 11 kilobase long genome codes a big polyprotein of more than 3,400 amino acids. This polyprotein is then cleaved into three structural proteins; core (C), membrane (M), and envelope (E) and seven non-structural ones, thus, NS1, NS2A, NS2B, NS3, NS4A, NS4B, and NS5. The construction of the virus particle is done by the structural proteins and the non-structural proteins are the most important proteins required in the viral replication and assembly inside the host cell. The capsid protein (C) is 120 amino acids long and it is important to enclose the viral RNA as well as to create the nucleocapsid core. The envelope (E) proteins and the precursor membrane (prM) proteins are glycoproteins; each of them is composed of two transmembrane domains. In the maturation of the virus, prM is cleaved to form pr peptide and M protein, and prM may possibly help the E proteins to correctly fold and assemble before being cleaved (10). The envelope (E) protein contains essential functional sites, among them, receptor-binding region and a fusion peptide, which help the virus to bind and enter host cells (11). All of these are categorized as dengue virus which is subdivided into four serotypes DEN-1, DEN-2, DEN-3, and DEN-4 according to their antigenic differences in the envelope proteins. These serotypes share similar structure, symptoms, and pathogenesis, though the factors influencing the severity of disease remain unclear (12).

**Fig 1.**
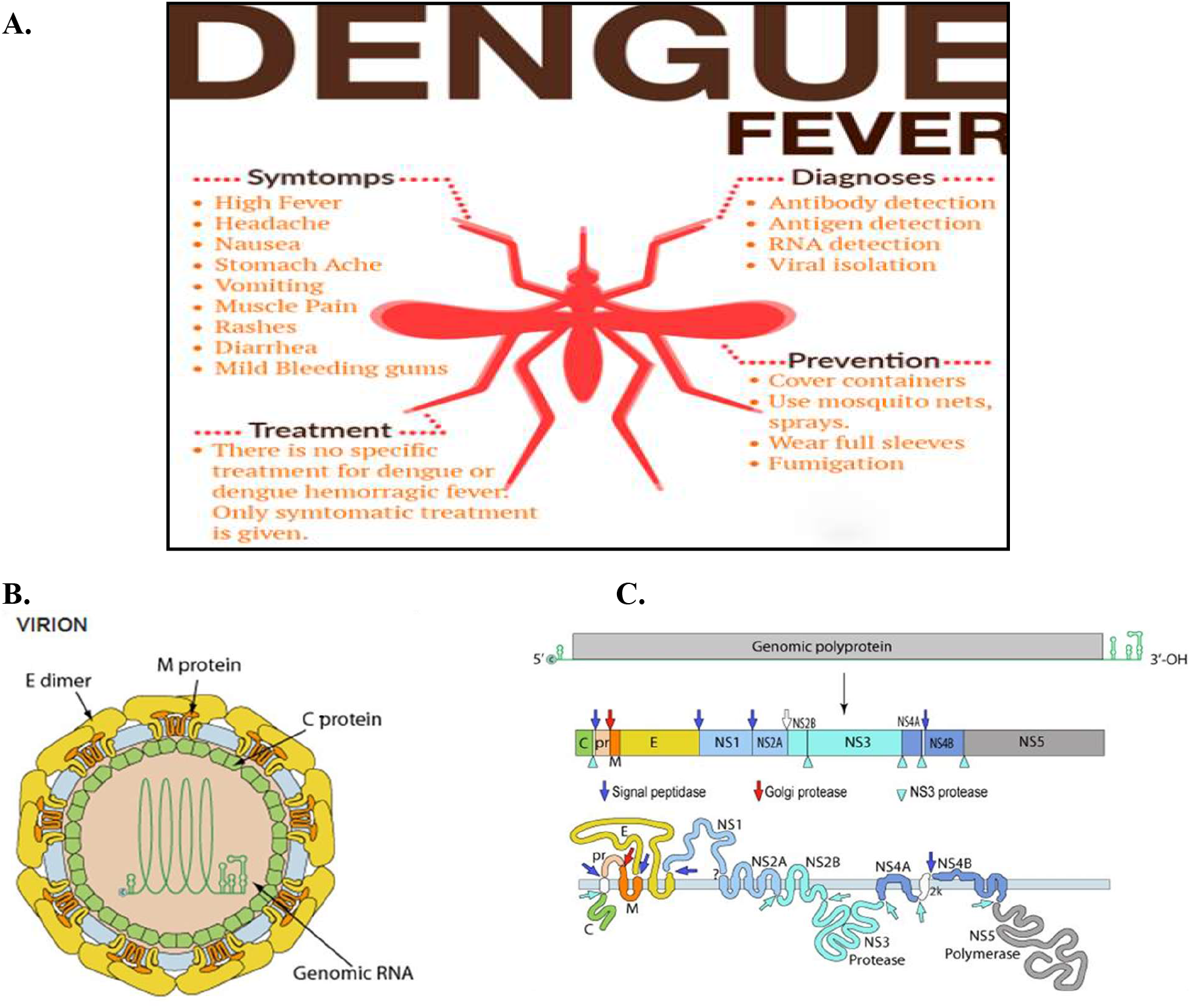
Depicts the structure and genome organization of the dengue virus. Panel (a) Overview of dengue fever (b) provides a schematic view of the structural proteins present in both immature and mature forms of the virus particles. Panel (c) illustrates the flavivirus genome organization and its expression pattern, highlighting the areas that code for structural and non-structural proteins, as well as the arrangement of the polyprotein within the membrane of the endoplasmic reticulum.

Dengue virus has a roughly spherical shape with an average diameter of about 50 nanometers. At its center lies the nucleocapsid, which includes the viral RNA genome tightly associated with capsid (C) proteins (13). The nucleocapsid core is enclosed by a lipid envelope that the virus obtains from the host cell as it replicates. In this lipid membrane, envelope (E) and membrane (M) proteins have more than 180 copies. These proteins form a protective outer coat which does not only protect the virus in addition to being essential in recognition and entry into host cells (14).

### 1.3. Dengue Viral Proteins and Its Function

Dengue virus precursors are three structural proteins that define the shape of the virus particle and seven nonstructural proteins that have significant roles in the replication and assembly of the viruses (15).

### 1.4. Structural Proteins

#### 1.4.1. Dengue Capsid Protein(C)

The capsid protein is a protein of such essential 12 kDa and is instrumental in the formation of the nucleocapsid as it interacts with the viral RNA. It is a dimer having a high net positive charge because the distribution of basic amino acids on its surface is not even (16). The nucleus localization of this protein is achieved by the presence of three nuclear localization signal at amino acid positions 6-9 (KKAR), 73-76 (KKSK), and a bipartite motif of RKRRRR at position 85-100. In the absence of these motifs, the protein of the capsid will live in the cytoplasm (17). Also, the signals allow communication with the apoptotic protein DAXX that has the potential to induce apoptosis. The primary purpose of the capsid protein is to secure the viral genome against environmental harm and help the delivery of viral genome to host cell. In the infected cells, it passes through a complicated two-step maturation procedure. The carboxyl terminus of the capsid protein acts as a membrane insertion sequence in translationary condition, through which a co-translational cleavage mediated by signals is performed, resulting in the release of the amino terminus of the prM protein (18). Before assembly of the virions, the C-terminal membrane-spanning fragment of the newly-produced intracellular capsid protein is broken off. It is possible that the viral NS3 protease can perform this cleavage in Trans, through the carboxyl-terminal side of various basic amino acid residues (19).

#### 1.4.2. Membrane Association Protein (M)

The membrane glycoprotein plays a critical role in shaping and maturation of virions and the form is built of seven antiparallel b-strands that are joined by three disulfide bonds (20). In a mature dengue virus, the outer shell has 180 copies of envelope (E) protein and membrane (M) protein that form 90 heterodimers in total, forming the ridges of the virus. The virus is exposed to acidic environments as the virus travels via secretory pathway -particularly trans-Golgi network (TGN). This low PH condition causes the envelope protein to dissociate with the prM protein and reform themselves into E protein homodimers, which lies flat to the viral surface to provide the maturing virus with a smoother appearance. This process of maturation continues through the dissociation of peptide with the M protein, although it remains attached to the E protein until the virus leaves the host cell (21).

#### 1.4.2. Envelope Protein (E)

Dengue virus envelope protein (E) is glycoprotein found on the outer side of the virus which plays the largest role during the attachment of the virus to the host cells. It is also receptive to different host cell receptors, some of which are Fc receptor, DC-SIGN, heat shock proteins HSP70/90, GRP78, Rab5, CD209, CD14, and the mannose receptor. Moreover, this protein targets heparin sulfate proteoglycans (HSPGs). The envelope protein has three domains, domain I (EDI), or a central structural domain that includes the N-terminus; domain II (EDII), or the dimerization domain that carries the site of attachment to the host cell by the envelope protein at its extreme end, and domain III (EDIII) or an immunoglobulin-like domain (23).

##### Non-Structural Proteins

###### Non-structural Protein 1 (NS1)

NS1 is a non-structural glycoprotein and has an approximate molecular weight of 50 kDa and it has significant roles during the viral life cycle and it is a major antigen target by anti-dengue antibodies (24). Once the viral RNA gets into the host cell, NS1 gets synthesized by the assistance of other non-structural proteins which are transcriptional regulators. When NS1 passes through endoplasmic reticulum (ER), it is glycosylated. There are three variants of NS1, which are located in different cellular compartments depending on their glycosylation (25). NS1 is first an immature monomer that is located in the ER and is critical in the replication of the virus. It may also be a stable homodimer that binds to membranes via a glycosyl-phosphatidylinositol (GPI) anchor (26). This membrane-enclosed form of NS1 is a possible binding site of NS1-specific antibodies and might mediate the signaling patterns of proteins of the GPI-type (27). The fully functional NS1 protein is 352 amino acids long and has 2 N -linked glycosylation sites, at 130 and 207, and 12 conserved cysteine residues-showing the significance of disulfide bridges to the structure and activity of NS1 in all flaviviruses. The dengue non-structural proteins, of which the NS1 is the only secreted protein, is thus unique because it is released as a soluble hexamer, and their scanners can be found in the patient serum on the first day of fever, and as late as three days later (28).

###### Non-structural Protein 2A

NS2A is a hydrophobic, membrane bound protein composed of 231 amino acids that carries with it a number of the most important functions especially during the replication of viral RNA. It can selectively bind to the untranslated region (UTR) of the viral RNA and bind to different elements of the replication complex. Also, NS2A assists in regulating antibody antiviral interferon reaction in the host and intervenes in the cycle of creation and discharge of virus particles (30). It is also involved in the delivery of new RNA synthesized to the location where the virus particles are formed. The indirect help of this RNA transportation could be provided by NS2A acting as a viroporin that creates holes in the membranes and helps to move (31).

###### Non-structural Protein 2B

NS2B serves as a companion to the serine protease associate of NS2B-NS3. This proteolytic core consists of the 40 last amino acids of NS2B and the 180 first of NS3 (32). This activates the protease complex by cleavage of the NS2B-NS3 precursor, which is essential in cleaving at the NS2A/NS2B junction. In NS2B, a stretch of 40 amino acids is essential in the activity of the proteases and includes a hydrophilic section that is enclosed by hydrophobic regions. It is detected that this hydrophilic part is critical to proteolytic activity (33). Moreover, the activity of type I interferon can be suppressed by the NS2B-NS3 protease complex, which inhibits the activity of the IFN-b promoter, the main mechanism of which is by blocking the phosphorylation of IRF3 (34).

###### Non-structural Protein 3

NS3 is the second-largest non-structural protein of the dengue virus and has a serine protease domain which is proximate to its N-terminal end with the positioning of NTPase, helicase and RNA triphosphatase (RTPase) domains which are at the C-terminal end. The protein plays a critical role in replication of the virus and is considered as a potential antiviral drug (35).

###### Non-structural Protein 4A

The C-terminal of NS4A protein is significantly hydrophilic and serves as a targeting sequence in its translocation to the lumen of the endoplasmic reticulum. It is a component of the membrane-bound viral replication complex the integrity of which is supported by NS4A (36). It plays a vital role in the process of virus replication, as it enables the cellular membranes to be remodeled and regulates the changes occurring. The N-terminal portion of NS4A also cleanses cell-free cleavage of a viral protease as was found in cell-free models (37).

###### Non-structural Protein 4B

It Is a small hydrophobic protein present in the association with the endoplasmic (38). It has the capability of controlling the replication of RNA with the aid of NS3. It is able to inhibit the innate immune system. NS4b is able to suppress the Tyk2 kinase activity by preventing the phosphorylation of the inhibition of STAT 1 even in the presence of IFN-B or IFN-y (39). It has been shown that NS4B presumably plays a role in suppressing interferon (IFN) signaling and is an IFN-signaling suppressor. Specifically, the dengue virus NS4B protein first 125 amino acids suffice to prevent the alpha/beta interferon (IFN-a/b) signaling pathway (40).

###### Non-structural Protein 5

Dengue virus serotype 5 (DEN5) protein NS5 protein contains 900 amino acids and has two functional domains. The first segment (amino acids 1-296), the N-terminal part, is a methyltransferase domain, and the second one (amino acids 320-900) is an RNA-dependent RNA polymerase (RdRp). NS5 is crucial to the viral RNA reproduction process primarily due to its RdRp activity that enables the virus to produce anew (de novo) RNA (41). Methyltransferase domain has a typical a/b/b sandwich, with both ends having sub domains at the N- and C-termini. The RdRp domain is similar to those of other RNA viruses, having palm, finger, and thumb subdomains as well as a preserved GDD motif that is necessary in piling on to the nucleotides during RNA translation (42).

**Table 2.**
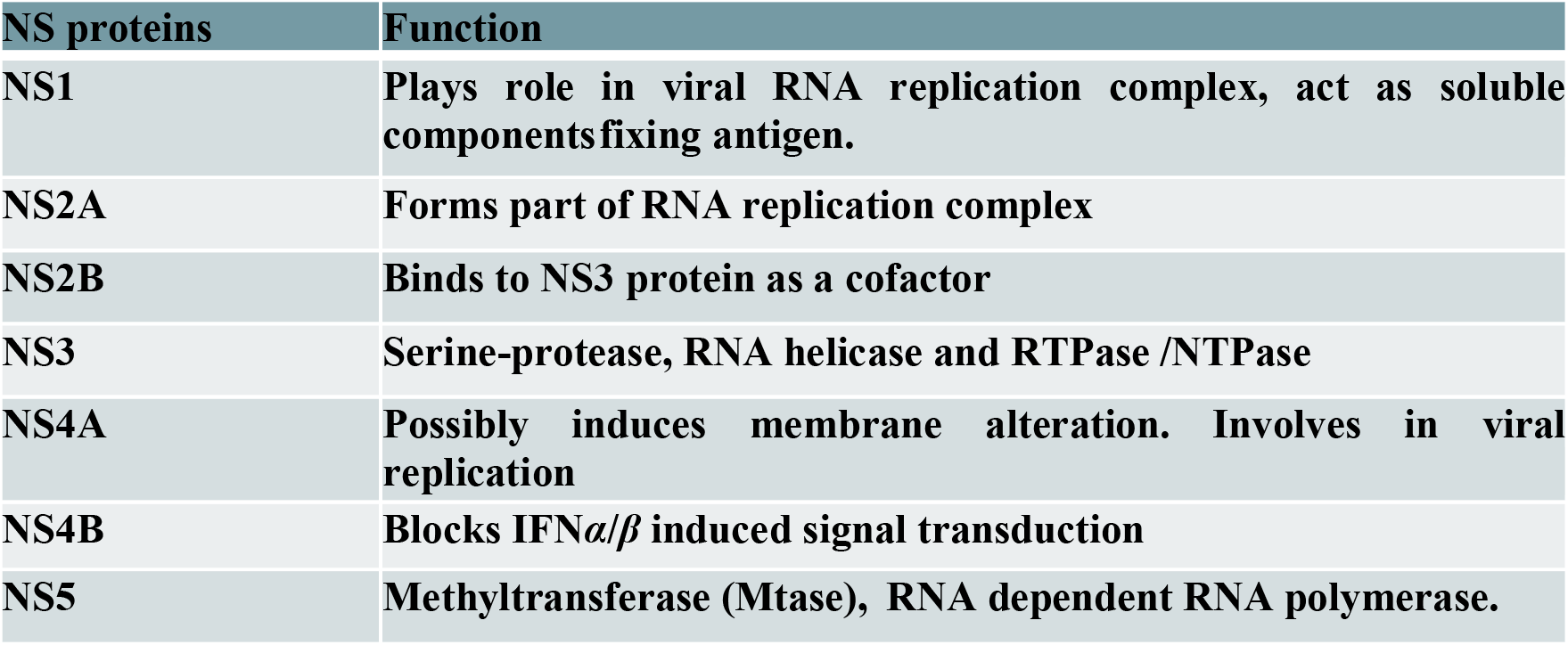
Functions of non-structural proteins.

| **NS proteins** | **Function** |
| --- | --- |
| **NS1** | **Plays role in viral RNA replication complex, act as soluble componentsfixing antigen.** |
| **NS2A** | **Forms part of RNA replication complex** |
| **NS2B** | **Binds to NS3 protein as a cofactor** |
| **NS3** | **Serine-protease, RNA helicase and RTPase /NTPase** |
| **NS4A** | **Possibly induces membrane alteration. Involves in viral replication** |
| **NS4B** | **Blocks IFN$\alpha/\beta$ induced signal transduction** |
| **NS5** | **Methyltransferase (Mtase), RNA dependent RNA polymerase.** |

###### Replication Mechanism

The dengue virus initiates infection by binding to specific receptors on the surface of the host cell and gaining entry via endocytosis. Following attachment, the virus merges with the endosomal membrane, which releases its contents into the cytoplasm. This process results in the disassembly of the viral particle and the liberation of its RNA genome. This viral RNA is then adopted into one long protein that is subsequently divided into ten viral proteins and the genome is also replicated. New viral particles are assembled on the endoplasmic reticulum, (ER) surface where structural proteins are integrated with new RNA that is recently synthesized. These ungrown viruses will then enter the trans-Golgi network (TGN) and become infectious forms. The developed dengue viruses eventually evade the host cell and infect other cells (45).

**Fig 2.**
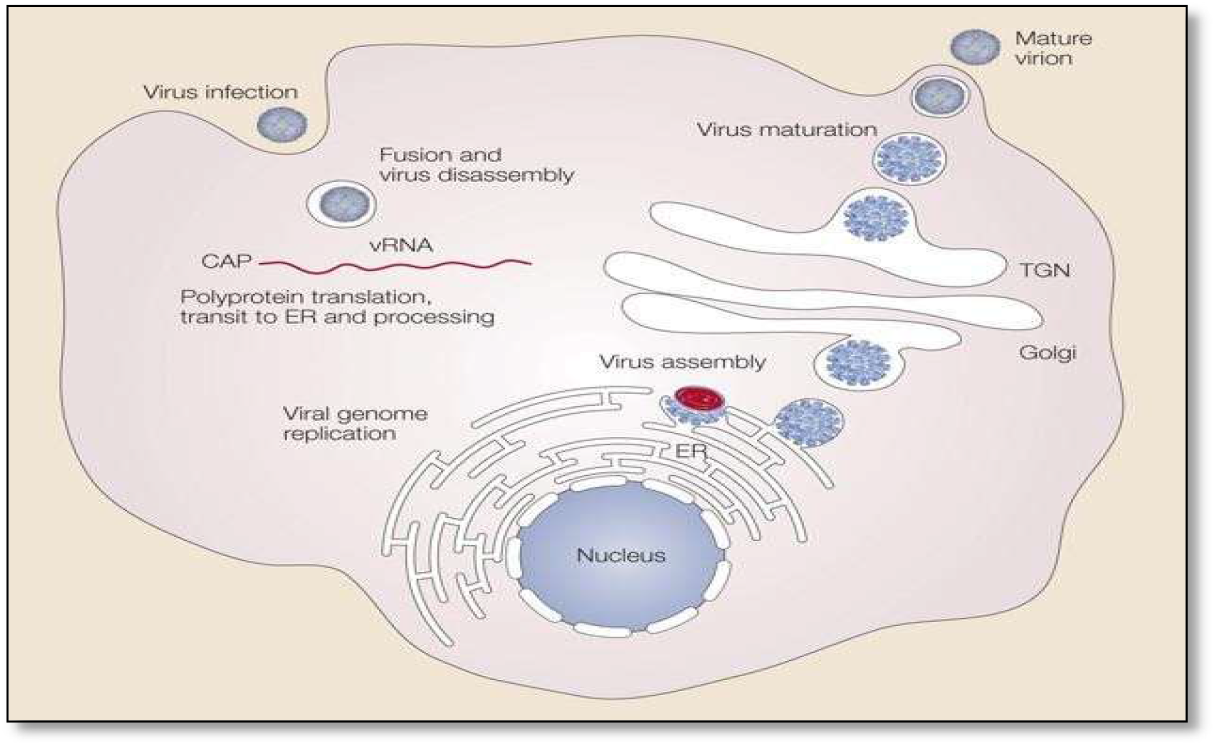
Demonstrates the replication mechanism of the dengue virus: the virus joins the surface receptors of the host cell and enters it through endocytosis. Having entered it combines with the endosomal membrane, and its RNA genome is discharged into cytoplasm. This RNA is then changed into one large polypeptide which becomes in its turn processed into ten different viral proteins as the genome is replicated. New viral particles are formed on the surface of the endoplasmic reticulum (ER) through the assembly of structural proteins and RNA. These immature viruses are then transported through the trans-Golgi network (TGN), where they mature into infectious forms. Ultimately, the mature viruses leave the host cell to infect additional cells.

###### Drugs Discovered To Inhibit Dengue Virus Replication

Bromocriptine (BRC)

ST-610

Benzoxazole

Posaconazole

Oxysterol-binding protein (OSBP)

###### Drugs and Its Modes of Action

Bromocriptine (BRC) has demonstrated strong anti-dengue virus (DV) effects while exhibiting low toxicity to cells. It works by blocking either the translation or replication phases of the DV life cycle (46). However, a single amino acid change (N374H) in the NS3 protein can cause resistance to BRC. Posaconazole is effective in suppressing the replication of both dengue and Zika viruses. Along with itraconazole, these antifungal medications inhibit ergosterol synthesis (47). Benzoxazole specifically targets the NS3 protein. Additionally, multiple dengue virus serotypes are now present in the same geographical areas (48).

###### Lauric Acid

Lauric acid or dodecanoic acid is a saturated fatty acid that has a 12-carbon chain, which categorizes it under medium-chain fatty acids. It is white powder that is light in color and has a slight scent that resembles bay oil or soap. Salts and lauric acid esters are called laurates. This acid is present in about half of the fatty acids of coconut milk, coconut oil, laurel oil, and palm kernel oil (not to be confused with palm oil) (49). It is present naturally in human breast milk (6.2 percent of total fat), in cow milk (2.9 percent) and goat milk (3.1 percent). Although lauric acid has the potential to raise the total cholesterol more than most of other fatty acids, this is principally because of an increase in the high-density lipoprotein (HDL), commonly referred to as good cholesterol. Findings have shown that lauric acid produces a better impact on increasing the total HDL cholesterol as compared to all other saturated or unsaturated fatty acids investigated. In the laboratory, lauric acid has found use in the determination of the molar mass of unknown compounds by freezing-point depression. It is suitable in this regard due to its high melting temperature of 43.8oC and cryoscopic constant of 3.9oC /kg/mol. The molar mass is determined through freezing of the mixture resulting after melting lauric acid with the unknown substance and measurement of the freezing temperature of the mixture (50).

###### BHK Cell Lines

BHK-21 (Baby Hamster Kidney Strain 21) is a fibroblast cell line that was obtained by culturing kidney tissues on baby hamsters. It has been extensively employed as the platform to undergo manipulation with the help of expression vectors containing selectable and amplifiable reporters (51). Large scale: BHK-21 cells are widely used in research studies that involve Dengue virus pathogenicity because they allow penetration and replication of Dengue virus. In addition, BHK-21 cells are susceptible to a number of other viruses such as human adenovirus D, retrovirus type 3, and vesicular stomatitis virus among others (52).

## MATERIALS AND METHODS

### Materials

The following resources were acquired for India’s Himedia Ltd.: DV2 replicon cells, geneticin (G418), foetal bovine serum (FBS), bovine serum albumin (BSA), Dulbecco’s Modified Eagle Medium (DMEM), Nunc tissue culture flasks, Alexa Fluor 488-conjugated anti-rabbit antibody from Thermo Scientific, a Leica DMI 3000 fluorescence microscope, Sigma’s lauric acid, MTT solution, a microplate reader, Renilla luciferase from Promega, and penicillin-streptomycin. Additionally, molecular grade dimethyl sulfoxide (DMSO) from Changzhou Yangquan Chemicals and ultrapure Milli-Q water from Merck-Millipore (Darmstadt, Germany) were procured from Sigma, India. All solutions and reagents were prepared using this ultrapure Milli-Q water.

### Methods

#### Naive Cells

Baby Hamster Kidney (BHK 21) cells served as the control (naïve) cells throughout this research. Both BHK-21 and DV2 replicon cells were sourced from the cell bank at the National Centre for Cell Science (NCCS) in Pune, India. These cells were grown in Dulbecco’s Modified Eagle Medium (DMEM) supplemented with 10% fetal bovine serum (FBS) and incubated at 37 °C in a humidified environment containing 5% CO_2_.

#### Maintenance of Stable DV2 replicon cells

The stable BHK 21 cells containing DV2-RNA were grown in DMEM medium supplemented with 10% fetal bovine serum (FBS) and incubated at 37◦C with 5% CO2. They were cultured under Genetic in (G418) antibiotic selection at a concentration of 400 μg/ml. The cells were regularly maintained by passaging every 3-4 days when they reached 60-70% confluence.

#### Morphological observation

Once the stable cells were generated, the morphology of the stable cells and Naive cells growing in a DMEM complete medium in a tissue culture flask (T-25 cm) were observed under bright filed microscope using 10 X magnification.

#### Detection of DV2-NS1 by Immuno fluorescence assay (IFA)

DMEM growth media containing 10 percent fetal bovine serum (FBS) and G418 was used to grow the stable cells on 1 cm2 glass coverslips, under normal conditions, 37oC, and 5 percent CO2. They were observed to grow after every 24 hours under phase-contrast microscope. The medium was disposed after 24 hours and the cells were fixed using 3.7% paraformaldehyde in phosphate-buffered saline (PBS) within 20 minutes at room temperature. The cells were then permeabilized using 0.1% Triton X-100 after which they were blocked using 5% horse serum after 15 minutes. After three PBS washes the cells were left overnight at 4 degC with a 1:1000 dilution of an anti-DV2-NS4A rabbit polyclonal antibody. The cells were washed thrice more with PBS and then incubated with the Alexa Fluor 488-conjugated anti-rabbit secondary antibody in the concentration of 1:200 over two hours in the room temperature. The staining solution existed of DAPI and lasted 10 minutes with three washes of PBS between them. Lastly, the coverslips were dried in the air and onto glass slides so that they could be examined under a fluorescence microscope. A Leica microscope (DMI 3000) was used to obtain immunofluorescence photomicrographs (20 X magnification) with the help of a camera.

#### The stable cells showed Luciferase activity

The stable cells harbouring DV2 replicon RNA were characterized for the presence of active transcription and translation of viral genes by assessing the Luciferase reporter activity in the presence and absence of MPA addition. The cell lysate obtained from the DV2-BHK21-replicon cells showed the luciferase activity of 96% higher as compared with the MPA treated stable BHK 21 cells (n=3 and p<0.0001). The decrease in the levels of luciferase relative luminescence units indicates, Mycophenolic Acid (MPA) addition inhibit the replication of DV2 replicon RNA thereby reduces the total viral protein translation along with luciferase reporter activity.

### Screening of Lauric acid against Dengue virus 2

#### MTT assay with auric Acid

Naive BHK-21 and DV2-BHK-Replicon cells were seeded in 96-well plates having a density of 10, 000 cells per well. Lauric acid was made in successive dilution to obtain the final concentrations of 5, 2.5, 0.625, 0.3125, 0.156, 0.0781, 0.0390, 0.0195, 0.0097 and 0.0048 uM. The cells experienced these levels of lauric acid and were allowed to incubate after 24 hours. Following incubation, 20 u L of MTT solutions was put into each well according to the instructions provided by the manufacturer. After 3 more hours of incubation, the medium was separated and 100 uL of DMSO was mixed into the dissolution of the resulting formazan crystals. The absorbance at 570 nm was then taken with the PerkinElmer multimode microplate reader. The percent cell viability at each concentration of lauric acid was computed and assessed by nonlinear regression with the help of GraphPad Prism v5 software.

#### Replicon Inhibition Assay with Lauric Acid

BHK-21 cells that had the DV2 replicon were grown in Dulbecco modified Eagle medium (DMEM) with 10% fetal bovine serum (FBS), 100 ug/mL penicillin-streptomycin, and 400 ug/mL G418. Cells of the 1×104 cell count in each well were plated up in a humidified chamber with 5% CO2 at 37 deg C and allowed to incubate after 24 hours. The cells were then subjected to lauric acid and left to incubate further after this period of 24 hours. The cells were then lysed and the Renilla luciferase (Rluc) activity was measured using Renilla Luciferase Assay System (Promega) according to the directions of the manufacturer. A PerkinElmer multimode plate reader was used to identify luminescence.

## RESULTS

### Morphological observation of BHK and DV2 Replicon Cells

**Fig 3.**
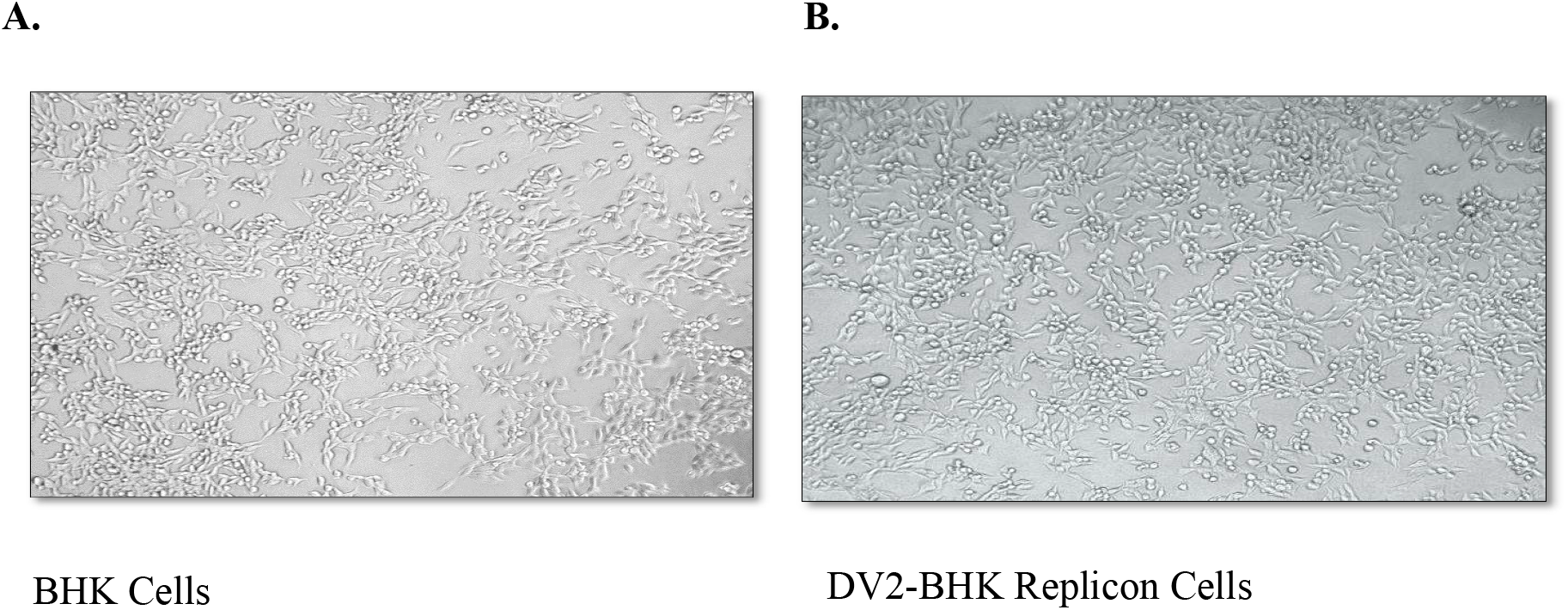
Morphological observation of cells in 10X magnification. Immunostaining of NS1 in DV2-BHK-Replicon cells.

**Fig 4.**
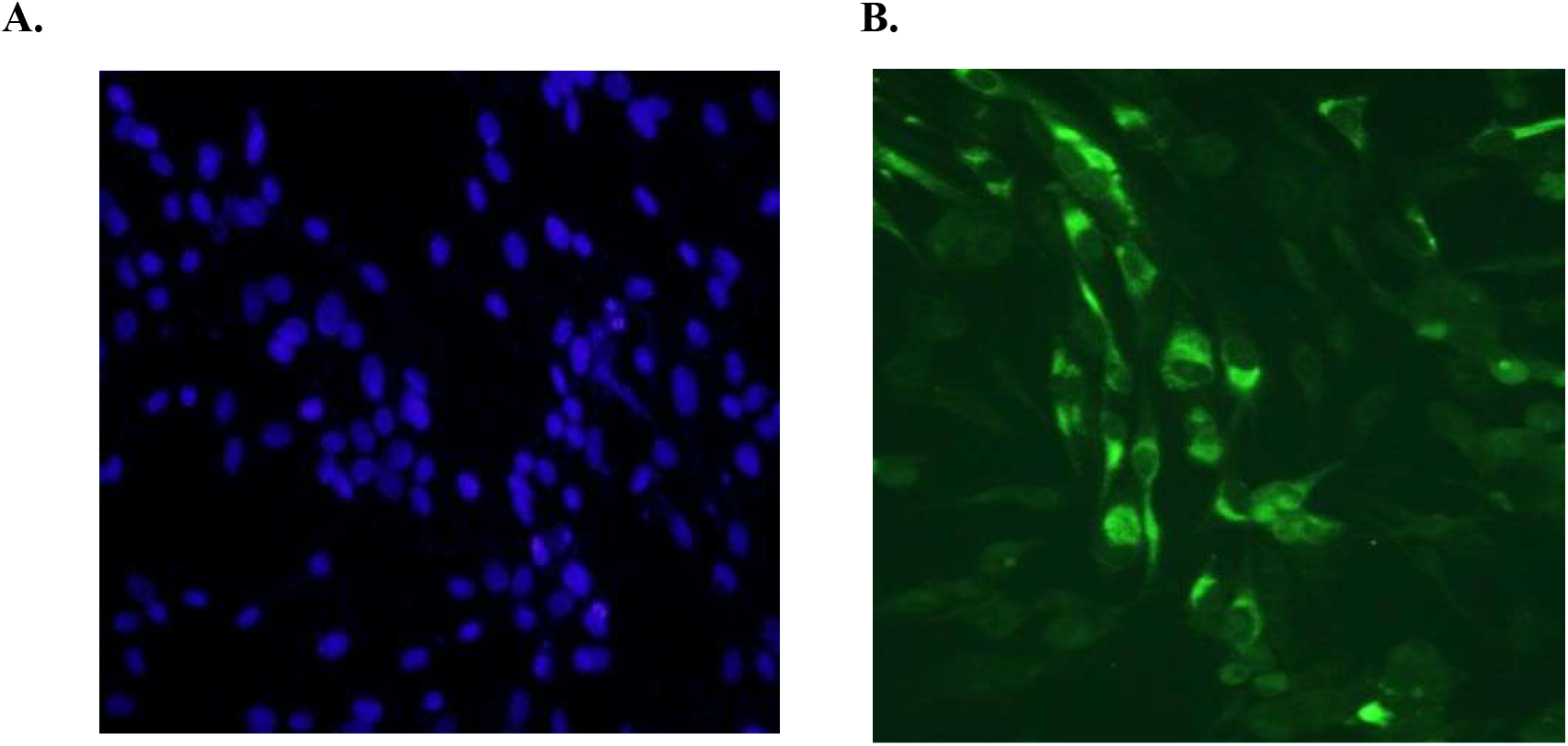
Immunostaining of NS1 in DV2-BHK-Replicon cells. (A) DV2-BHK-Replicon cells stained with DAPI. (B)Immunostained with Anti-NS1 antibody.

### Luciferase Activity

**Fig 5.**
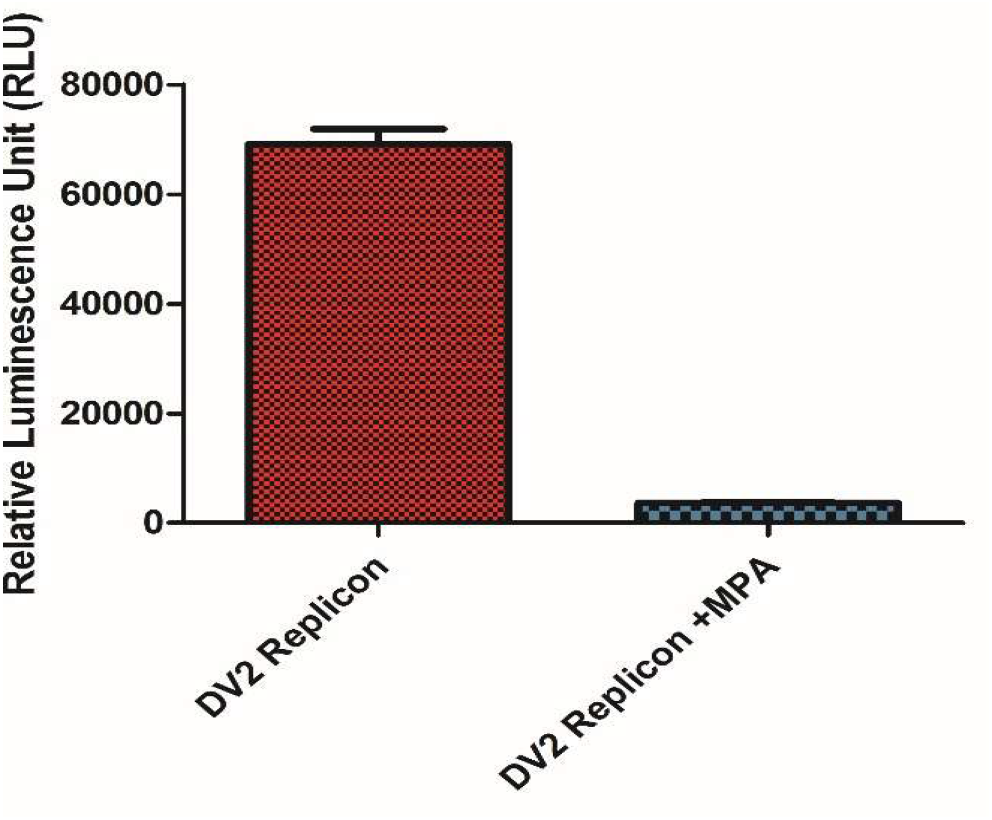
Luciferase Activity: The decrease in the levels of luciferase relative luminescence units indicates, Mycophenolic Acid (MPA) addition inhibit the replication of DV2 replicon RNA thereby reduces the total viral protein translation along with luciferase reporter activity. MPA has been used as a positive control in this study.

### Cytotoxicity assay of Lauric acid

**Fig 6.**
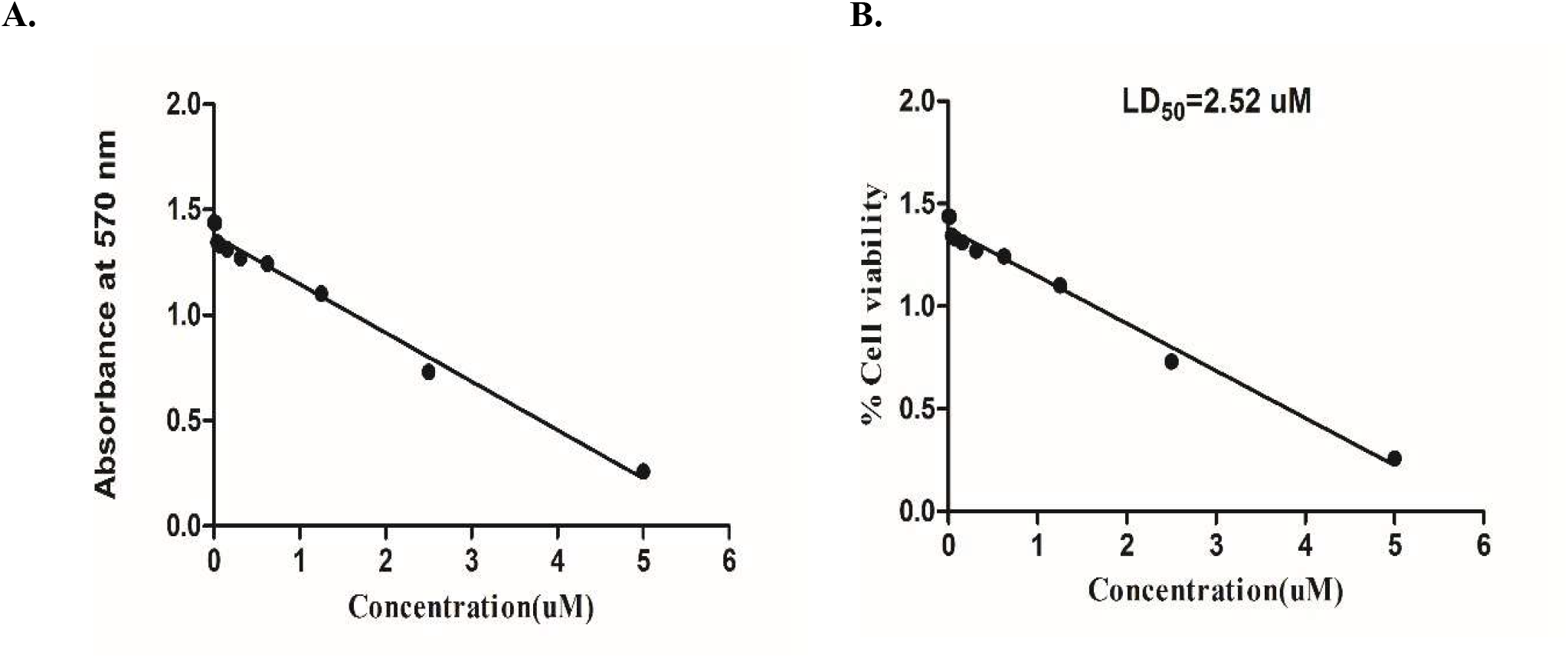
illustrates the cytotoxicity test of Lauric Acid on DV2 Replicon Cells. Lauric acid was diluted in a series to achieve final concentrations of 5, 2.5, 0.625, 0.3125, 0.156, 0.0781, 0.0390, 0.0195, 0.0097, and 0.0048 µM for the evaluation. (A) Concentration versus Absorbance at 570 nm: An MTT assay was conducted 24 hours post-treatment, measuring absorbance at 570 nm to assess cellular metabolic activity. (B) Cell Viability Curve: The percentage of viable cells was determined by the formula: (Absorbance of treated cells / Absorbance of control cells) × 100. Both the half-maximal inhibitory concentration (IC_50_) and the lethal dose 50 (LD_50_) were determined to be 2.52 µM, marking the cytotoxic limit of Lauric Acid. (C) Morphological Changes: Panel A displays DV2 replicon cells immediately following Lauric Acid treatment, while Panel B presents the same cells 24 hours later, highlighting morphological changes resulting from cytotoxicity.

### Cytotoxic effect of Lauric acid in Dengue virus 2 replicon cells

When Lauric acid was applied at elevated levels (5 µM, 2.5 µM, and 1.25 µM), a significant cytopathic effect (CPE) appeared in DV2 replicon cells. The monolayer displayed severe disruption, characterized by rapid cell shrinkage, increased cell density, and complete detachment from the surface of the culture flask within 72 hours post-treatment. This form of CPE represents the most severe cellular damage and is typical of flavivirus-infected cells under cytotoxic stress.

**Fig 7.**
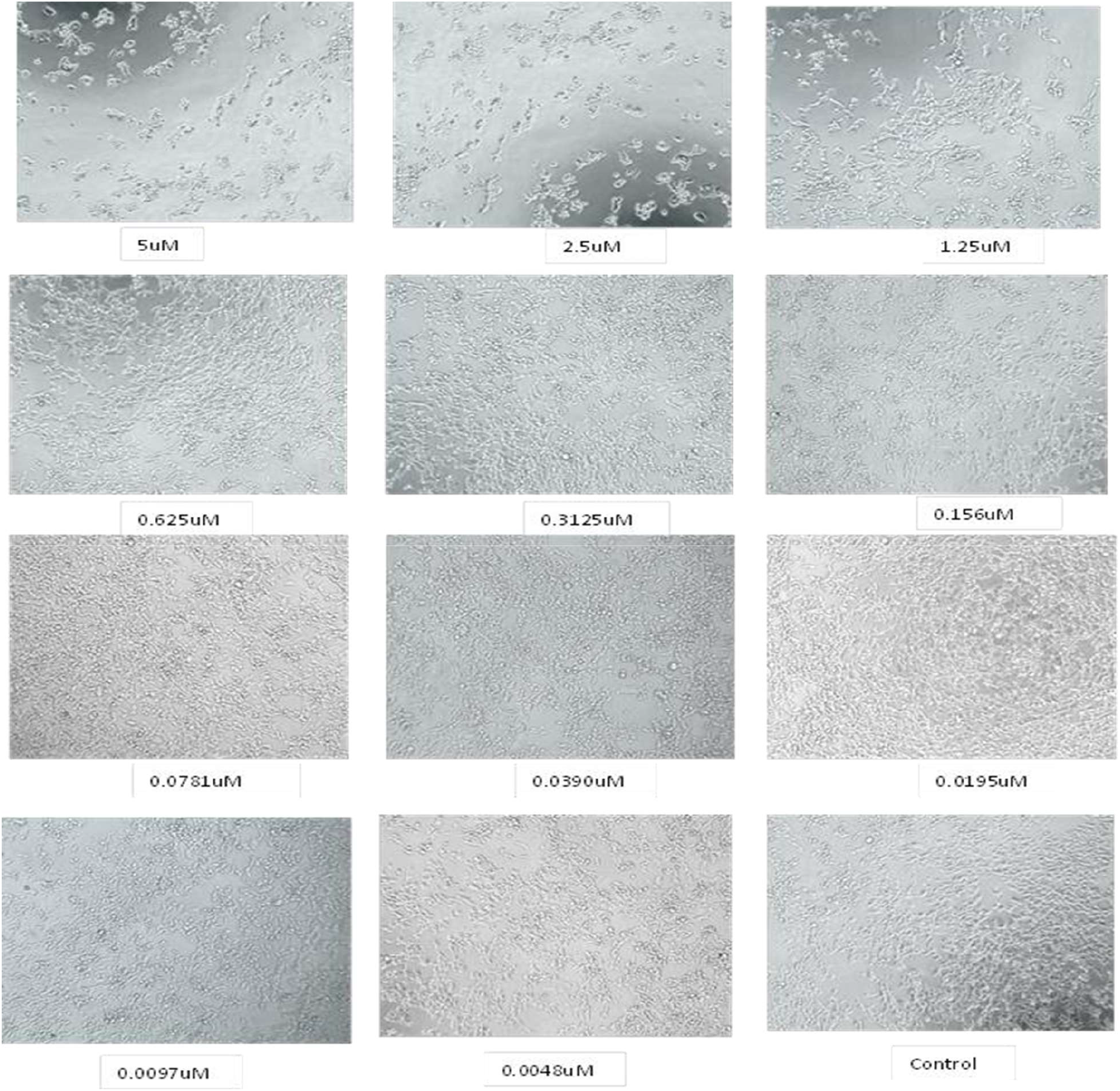
Cytotoxic effect of cells. upon treatment of Lauric acid in different concentrations (5,2.5, 0.625, 0.3125, 0.156, 0.0781, 0.0390, 0.0195, 0.0097, 0.0048 µM).

### 3.6 Replicon Inhibition Assay

**Fig 8.**
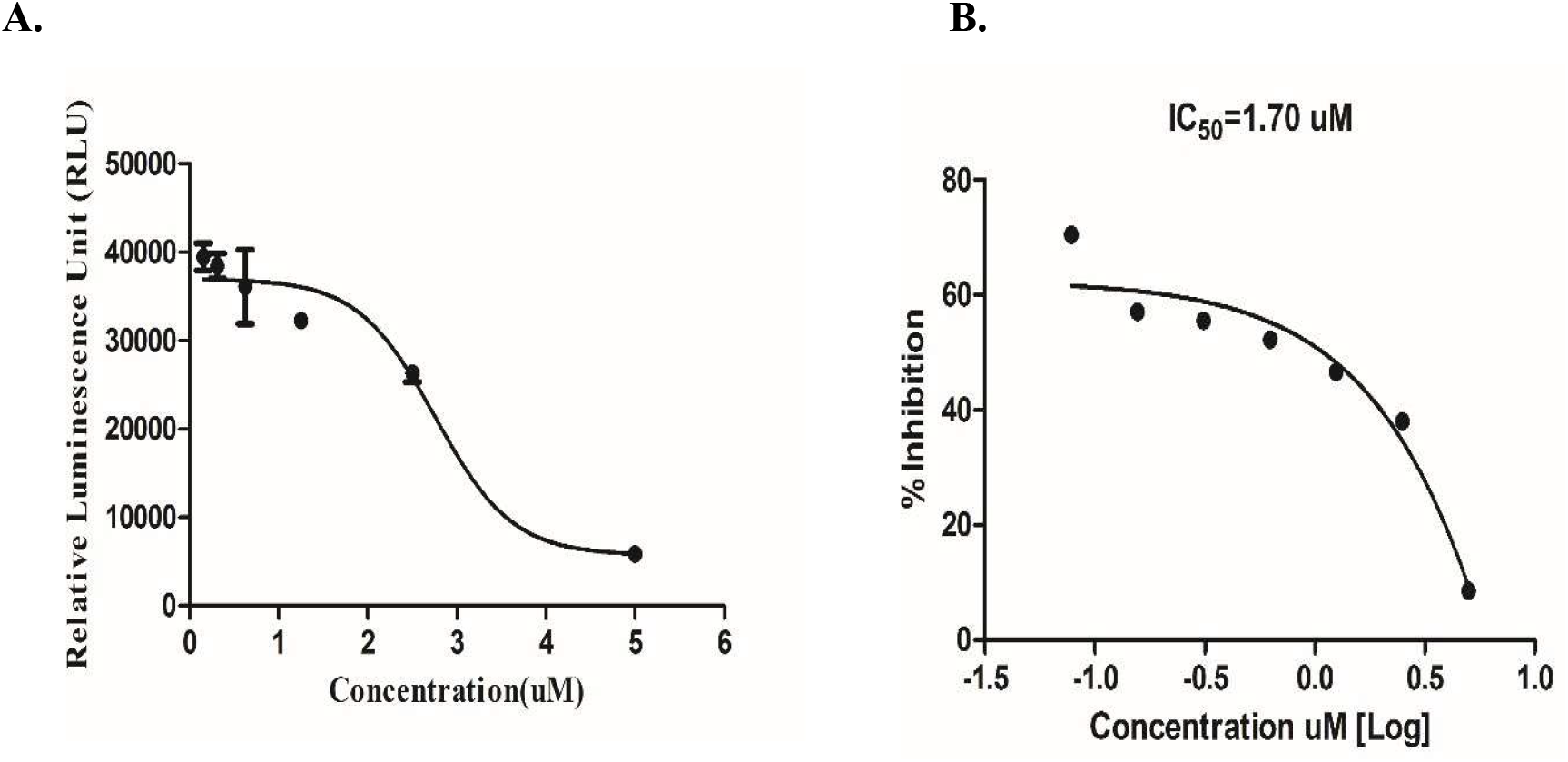
Replicon Inhibition Assay with Lauric Acid: BHK-21 cells expressing DV2-replicon, were seeded into 96-well micro plate and treated with Lauric acid and incubated for 24 hours with various concentrations (5, 2.5, 0.625, 0.3125, 0.156, 0.0781, 0.0390, 0.0195, 0.0097, 0.0048 µM). luciferase (Rluc) activities were measured and Inhibition Concentration 50 (IC50) was calculated. (i) Relative Luminescence unit (RLU) Vs Concentration. (ii)Percentage of inhibition was calculated and found to be IC50=1.70 uM.

## DISCUSSION

Dengue is a global health concern which affects two third of the global population. The disease induces significant morbidity and mortality and results in the severe economic impact in the growing country like India. However, there are no clinically approved drugs or vaccine against the dengue viruses (53). Dengue viruses are classified into four serotypes that are antigenically different, known as DV1 through DV4. It has been reported that dengue virus exploits host cell lipid metabolism for the successful replication along with the other host cell as well as viral components (54). There are many drugs, which are reported to be inhibitors against dengue virus protease (NS3) and RNA dependent RNA polymerase (NS5). However, targeting of drugs against these viral proteins may induce drug resistance. Hence, this study aimed to examine the impact of the small-chain fatty acid lauric acid, which has a 12-carbon chain, on the replication of the dengue virus (55). Lauric acid is the primary component of virgin coconut oil and has long been recognized for its antimicrobial properties. The study was conducted in stable BHK21 cells harboring dengue virus serotype 2 sub genome, expressing all nonstructural proteins (NS1 to NS5) along with the luciferase reporter gene. The genome replication and translation were characterized by the expression analysis of NS1 immunostaining (56). Lauric acid is found to be toxic to the replicon cells and LD 50 value was 2.52 uM, as evidenced by the MTT assay. Meantime, the drug was inhibiting the dengue virus 2 genome as evidenced by the reduction in the luciferase activity (IC50 = 1.50 uM) (57). This dose was closer to the IC50 value of mycophenolic acid (1.25 uM), known inhibitor of dengue virus replication (Diamond M et al, 2000). Therefore, the study indicates that small chain fatty acid such as lauric acid can be used as potential dengue virus inhibitor. The mechanism of action of these fatty acids need be explored at molecular level. Further, the toxicity of lauric acid needs to be mitigated by proper drug delivery platform.

## ACKNOWLEDGMENTS

The Department of Biotechnology and Department of Botany at Vinoba Bhave University, Hazaribagh, Jharkhand, is acknowledged by the authors for providing the entire infrastructure required to conduct the study.

## AUTHOR CONTRIBUTIONS

Anjali Kumari, investigation, and wrote original draft, edited, and supervised the research | Rajendra Pilankatta, conceptualization, funding acquisition, methodology| Bharti Kumari, Formal analysis, Investigation| Mihir Kumar Prasad, Formal analysis, Methodology | Nikhil Kumar, conceptualization and investigation | Anubha Kumari, methodology, edited, collected articles and shared in writing of original draft. All authors reviewed the final manuscript and agreed to the published version of the manuscript.

## DATA AVAILABILITY

Data will be made available on request.

